# Investigating the Role of Chromatin Remodeler FOXA1 in Ferroptotic Cell Death

**DOI:** 10.1101/2021.10.13.461056

**Authors:** Emilie Logie, Louis Maes, Joris Van Meenen, Peter De Rijk, Mojca Strazisar, Geert Joris, Bart Cuypers, Kris Laukens, Wim Vanden Berghe

## Abstract

Ferroptosis is a lipid peroxidation-dependent mechanism of regulated cell death known to suppress tumor proliferation and progression. Although several genetic and protein hallmarks have been identified in ferroptotic cell death, it remains challenging to fully characterize ferroptosis signaling pathways and to find suitable biomarkers. Moreover, changes taking place in the epigenome of ferroptotic cells remain poorly studied. In this context, we aimed to investigate the role of chromatin remodeler forkhead box protein A1 (FOXA1) in RSL3-treated multiple myeloma cells because, similar to ferroptosis, this transcription factor has been associated with changes in the lipid metabolism, DNA damage, and epithelial-to-mesenchymal transition (EMT). RNA sequencing and Western blot analysis revealed that FOXA1 expression is consistently upregulated upon ferroptosis induction in different *in vitro* and *in vivo* disease models. *In silico* motif analysis and transcription factor enrichment analysis further suggested that ferroptosis-mediated FOXA1 expression is orchestrated by specificity protein 1 (Sp1), a transcription factor known to be influenced by lipid peroxidation. Remarkably, FOXA1 upregulation in ferroptotic myeloma cells did not alter hormone signaling or EMT, two key downstream signaling pathways of FOXA1. CUT&RUN genome-wide transcriptional binding site profiling showed that GPX4-inhibition by RSL3 triggered loss of binding of FOXA1 to pericentromeric regions in multiple myeloma cells, suggesting that this transcription factor is possibly involved in genomic instability, DNA damage, or cellular senescence under ferroptotic conditions.

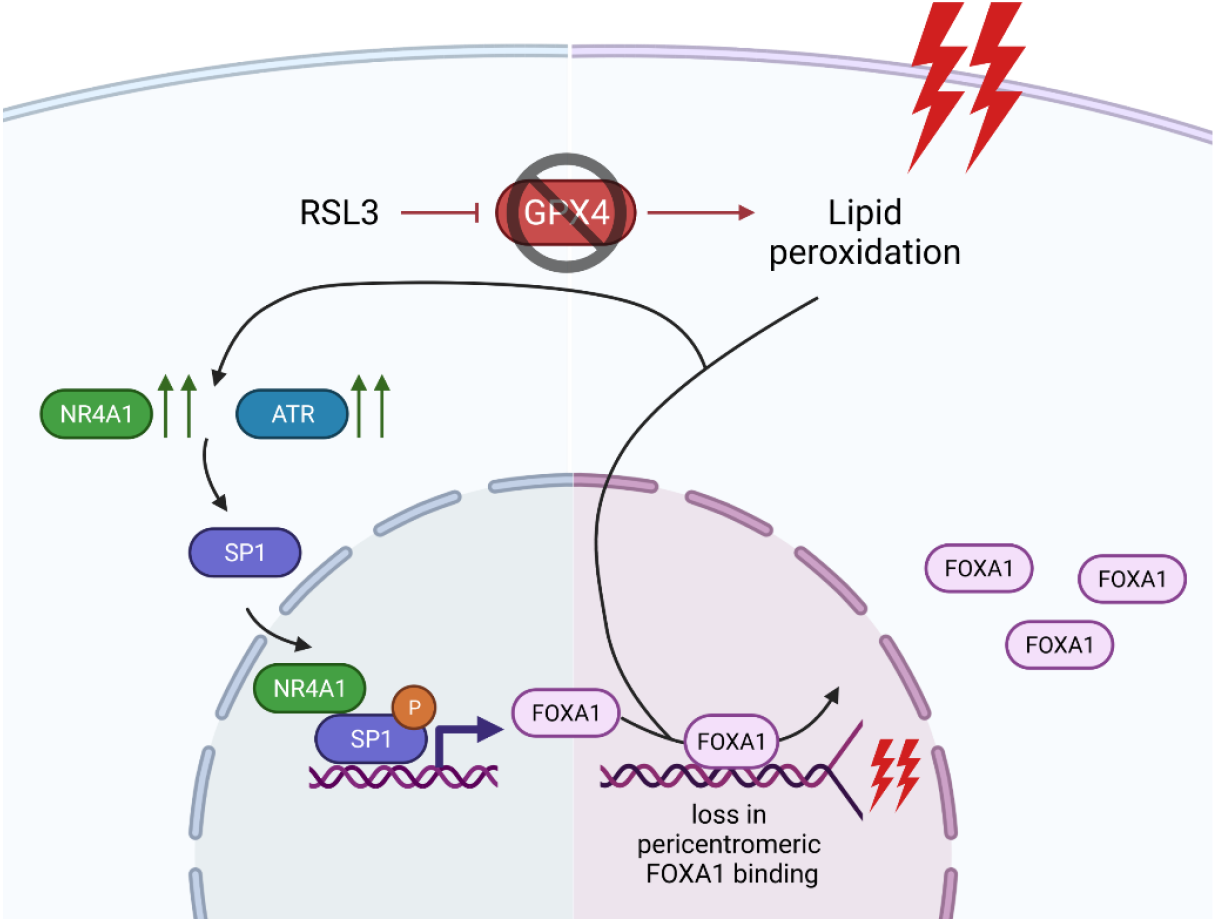

## Introduction

Ferroptosis is a non-apoptotic mode of regulated cell death (RCD) characterized by an iron-dependent rise in reactive oxygen species (ROS) that propagate lipid peroxidation reactions [1]. Mechanistically, intracellular increases in labile ferrous iron (Fe^2+^) challenge the cellular anti-oxidant defense systems by triggering the formation of toxic hydroxyl radicals through Fenton and Fenton-like chemistry [2]. Should enzymatic anti-oxidants, such as superoxide dismutase or glutathione peroxidases, fail to eliminate these toxic by-products, excessive peroxidation of polyunsaturated fatty acids (PUFAs) will ensue and ultimately cause detrimental loss of membrane integrity [3]. Several pathologies, including neurodegenerative diseases, cardiovascular diseases, ischemia-reperfusion injuries, and diabetes, have already been associated with ferroptotic cell death [4-8]. Small molecules targeting ferroptosis signaling pathways have therefore gained considerable clinical interest in the past couple of years [9]. Interestingly, the induction of ferroptosis has also demonstrated to offer therapeutic potential, especially in the field of oncology. One of the major hallmarks of cancer cells includes evasion of apoptotic cell death due to (acquired) therapy resistance mechanisms [10]. Provoking non-apoptotic modes of cell death, such as ferroptosis or necroptosis, might therefore help in eliminating therapy-resistant cancer (stem) cells [11]. Additionally, compared to healthy tissue, malignant tumors heavily rely on an increased iron metabolism to sustain their augmented proliferation capacity, exposing them to higher basal levels of oxidative stress [12]. Further elevating intracellular Fe^2+^ concentrations with ferroptotic compounds might further disturb their precarious redox balance and efficiently promote cell death [13]. For example, several B-cell malignancies, including multiple myeloma (MM) and B-cell lymphomas, portray an increased iron uptake and display sensitivity to ferroptosis inducers [14-20].

On a molecular level, genetic and protein hallmarks of ferroptosis have been identified and are mainly involved in oxidative stress pathways (NRF2, GPX4, CHAC1) [17, 21, 22], iron metabolism (TFRC, FTH1) [23, 24], inflammation (PTGS2) [17], and lipid metabolism (ACSL4) [25]. The overexpression or downregulation of these genes have been considered as potential biomarkers of ferroptosis cell death, yet it remains challenging to find ferroptosis-specific markers [26]. ACSL4, for instance, is currently considered to be a specific driver for ferroptotic cell death as it is involved in enhancing PUFA content in phospholipid bilayers, which are most susceptible to lipid peroxidation [25, 27]. However, a recent study by Chu and colleagues has demonstrated that even ACSL4-depleted cells can undergo p53-mediated ferroptosis [28]. Thus, there is an unmet need for finding more precise and specific contributors of ferroptotic cell death. A (combination of) suitable ferroptosis biomarker(s) might not only offer new insights in designing novel therapies for iron-related diseases, but might also aid in early detection of ferroptotic cells [29]. Moreover, it could help identify ferroptosis-resistant cancers, which, unfortunately, have already been identified as well [30-33].

In the present study, we investigated the role of chromatin remodeler forkhead box A1 (FOXA1) in MM cells undergoing ferroptotic cell death. FOXA1 belongs to a large family of FOX pioneer TFs that, unlike most TFs, can access target sequences located on nucleosomes and on some forms of compacted chromatin [34]. It is believed that members of the FOXA subfamily stably bind to genomic regions prior to activation and prior to binding of other TFs, and promote ATP-independent chromatin opening to allow binding of other TFs, nucleosome remodelers, or chromatin modifiers [34]. In case of FOXA1, it is suggested that chromatin opening is promoted by simultaneous DNA- and core histone binding (through a C-terminal domain), which disrupts local internucleosomal interactions required for stability of higher-order chromatin structure [35]. Depending on its chromatin recruitment sites, FOXA1 plays a role in embryonic development [35], hormone regulation [36, 37], lipid metabolism [38, 39], epithelial-to-mesenchymal transition (EMT) [40, 41], and DNA damage [42]. Given that the three latter processes have directly been linked to ferroptosis sensitivity or ferroptotic cell death [27, 43, 44], we combined RNA and CUT&RUN sequencing to characterize FOXA1 expression profiles and downstream targets in different ferroptosis models.

## Results

### Ferroptotic Cell Death Promotes FOXA1 expression in Different Disease Models

Although inhibition and induction of ferroptotic cell death is extensively being studied as a therapeutic strategy in several disease models, finding suitable ferroptosis biomarkers remains challenging [26]. To identify key genetic hallmarks of ferroptosis signaling pathways, we compared publicly available RNAseq data (GSE104462) of erastin-treated HEPG2 liver cancer cells to our own RNAseq data of RSL3-treated MM1 myeloma cancer cells (awaiting GEO accession number). Despite considerable differences in experimental design and starting material (Table 1), we found 23 common significant (FDR < 0.05 & │log2FC > 1│) differentially expressed genes (DEGs) that displayed a similar pattern in gene expression upon ferroptosis induction (Figure 1a). These genes are mainly involved in metal binding (YPEL5, ZBTB10, MT2A, MT1F, MT1X), DNA binding (BHLHE41, FOXA1, MAFF, KLF2, NR4A2), protein ubiquitination (HERPUD, PELI1, FBXO32), calcium ion binding (STX11, JAG1), and protein dephosphorylation (DUSP4, DUSP5). Interestingly, we could identify the ATP-independent chromatin remodeler FOXA1 as one of the common genes between both RNAseq datasets. FOXA1 is a 473 amino acid long TF that belongs to the family of FOX pioneer TFs. Through its winged forkhead domain (FKHD), it is able to open chromatin by disrupting internucleosomal interactions (Figure 1b). In agreement with the RNAseq data, qPCR and Western blot analysis revealed a time-dependent upregulation of FOXA1 expression in therapy-resistant and – sensitive MM1 cells treated with RSL3, a class II ferroptosis inducer (Figure 2a-c). As prolonged treatment with RSL3 results in decreased cell viability, these data suggest that FOXA1 expression is tied to severity of ferroptotic cell death and GPX4 inhibition (Figure 2b).

**Table 1:** Overview of experimental design differences in public RNAseq data vs own RNAseq data

| Feature | Public RNAseq data<br>(GSE104462) | Our RNAseq data |
| --- | --- | --- |
| Cell line | HEPG2 | MM1S & MM1R |
| Tissue of origin | Liver | Peripheral blood |
| Ferroptosis inducer | 10 μM erastin<br>(inhibits Xc<sup>-</sup> system) | 5 μM RSL3<br>(inhibits GPX4) |
| Duration ferroptosis treatment | 24 hr | 3 hr |
| Ferroptosis inhibitor | 1 μM ferrostatin-1 | 2 μM ferrostatin-1 |

Abbreviations: Xc<sup>-</sup> system, Cystine/glutamate transporter; GPX4, Glutathione peroxidase 4.

**Figure 1:**
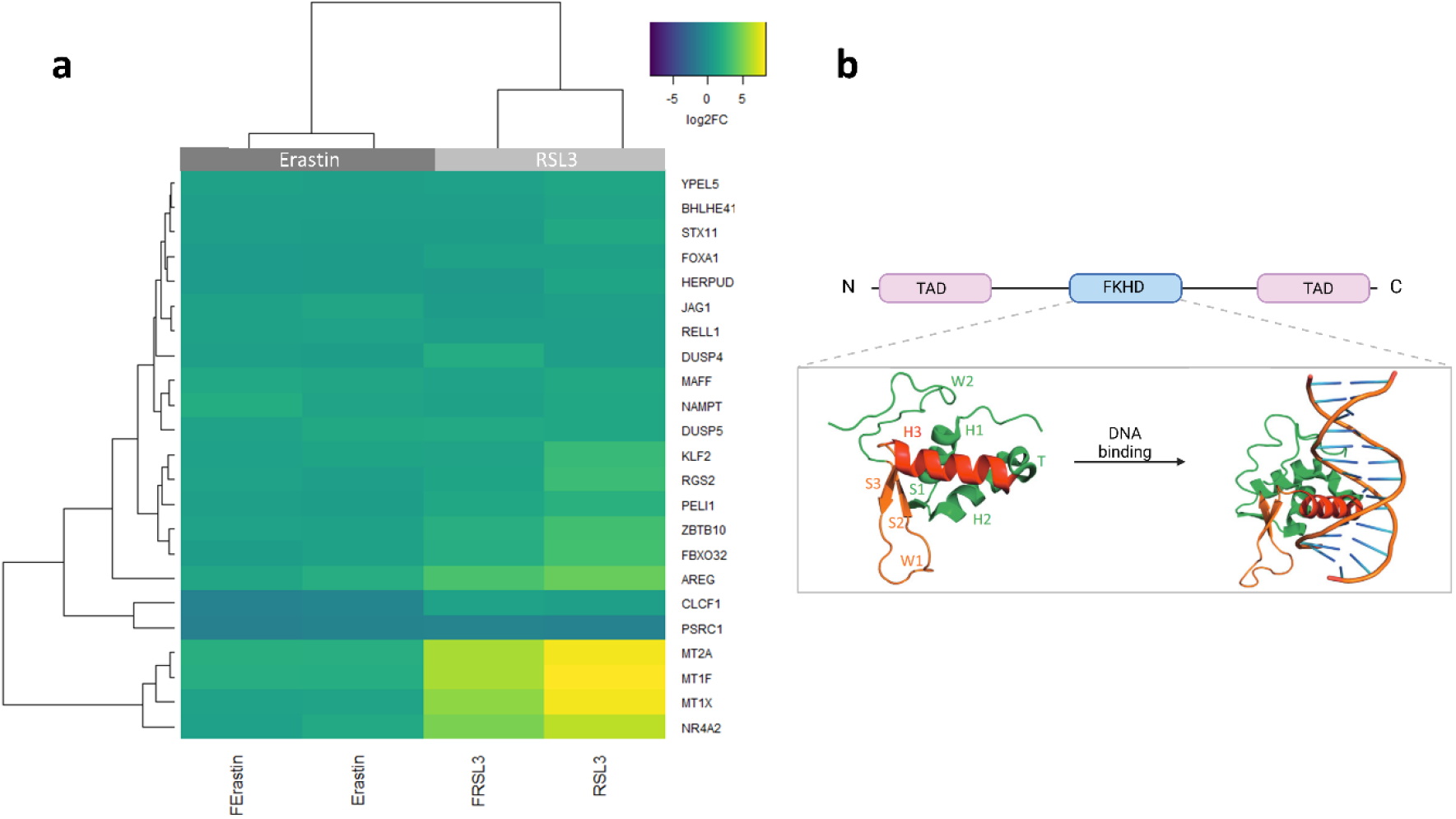
(**a**) Heatmap representation of common differentially expressed genes (FDR < 0.05, logFC > | 1 |) between erastin-treated HEPG2 cells (publicly available data GSE104462) and RSL3-treated MM1 cells. *N*=3 biologically independent replicates per cell line. (**b**) Schematic overview of Forkhead box A1 (FOXA1) protein domains. The forkhead domain (FKHD) is crucial for DNA binding and consists of 3 α-helices (H1-3) and 3 β-sheets (S1-3) organized in a helix-turn-helix motif. This motif is flanked on both sides by polypeptide chain “wings” (W1-2) that interact with the minor DNA groove.

**Figure 2:**
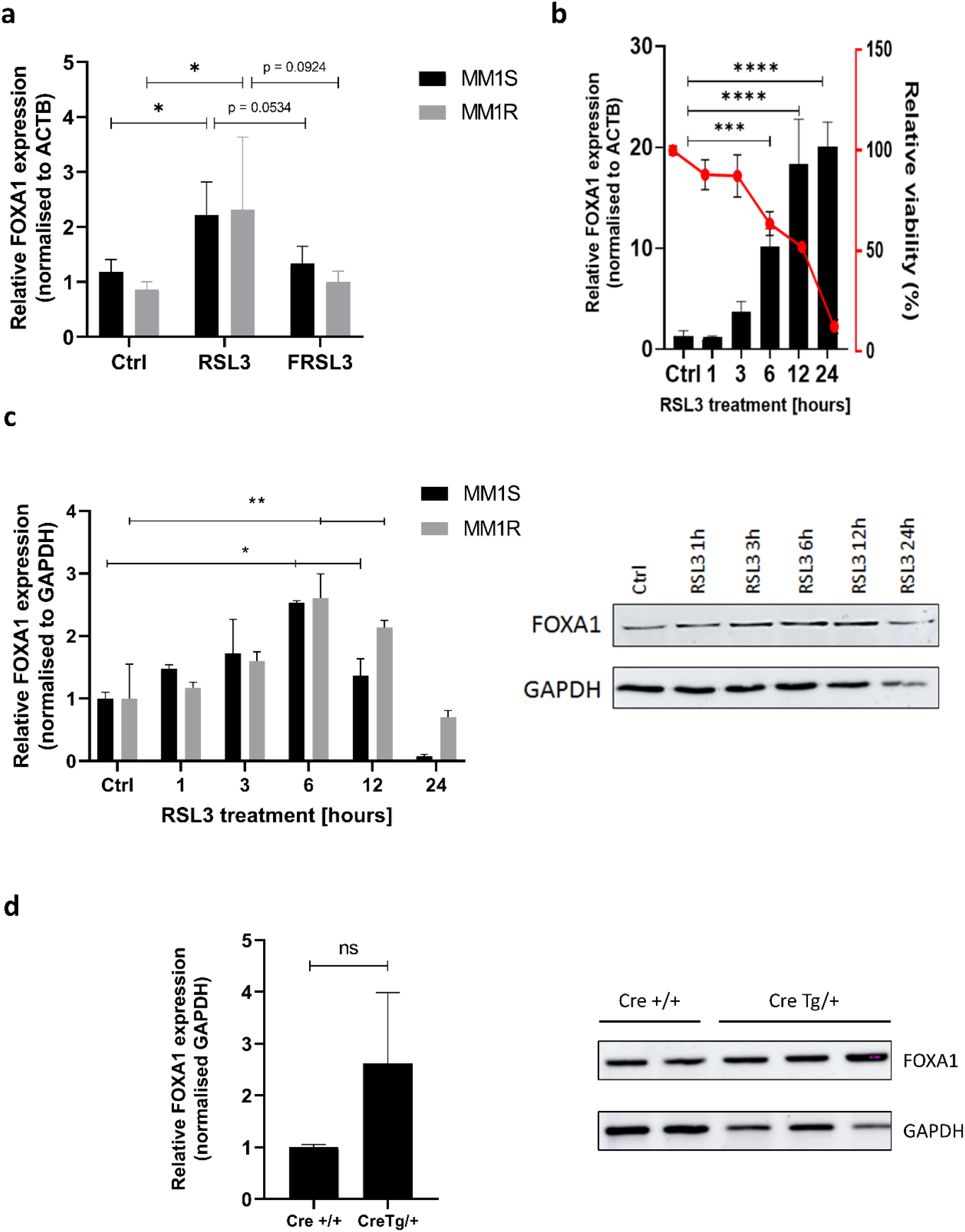
(**a**) Relative FOXA1 mRNA expression in MM1R and MM1S cells treated with 5 µM RSL3 for 3 hrs with (FRSL3) or without (RSL3) 2 hr pre-treatment with 2 µM ferrostatin-1 compared to untreated controls. FOXA1 expression is normalized against the β-actin (ACTB) housekeeping gene. Data are plotted as the mean ± s.d., *n*=3 biologically independent samples per cell line (*p < 0.05), ANOVA). (**b**) Relative mRNA FOXA1 expression and cell viability (%) in MM1 cells after RSL3 treatment. FOXA1 expression is normalized against ACTB mRNA expression. Data are plotted as the mean ± s.d., *n*=3 biologically independent samples per cell line (***p < 0.001, ****p < 0.0001, ANOVA). (**c**) Western blot detection and quantification of FOXA1 and GAPDH expression levels in MM1 cells treated with RSL3. Data are plotted as the mean ± s.d., *n=*3 biologically independent samples. (**d**) Western blot detection and quantification of FOXA1 and GAPDH expression levels in liver samples from healthy Cre +/+ mice versus sick Cre Tg/+ mice. Data are plotted as the mean ± s.d., *n*= 2 Cre +/+ mice and 3 Cre Tg/+ mice (ns = p > 0.05, two-tailed t-test).

Given that we detected ferroptosis-mediated FOXA1 induction in two different cell types (i.e. MM1 and HEPG2), we questioned whether similar observations could be made in *in vivo* ferroptosis models. To this end, we performed Western blot analysis on liver samples isolated from GPX4 liver-specific inducible knockout mice (Supplementary Figure S1). LoxP-GPX4 homozygous mice carrying the *cre* transgene (Cre Tg/+) demonstrated an increased, yet not significant, FOXA1 protein expression compared to their healthy controls (Cre +/+) (Figure 2d). Taken together, these findings suggest that FOXA1 upregulation may be a universal phenomenon in different ferroptotic (disease) models and that FOXA1 might be a central regulator in ferroptosis signaling.

### FOXA1 Binding to Pericentromeric DNA Regions is Reduced Under Ferroptotic Conditions

A PubMed search of all articles featuring the FOXA1 transcription factor revealed that FOXA1 expression is mostly associated with hormone signaling in prostate, breast and testis cancer (Supplementary Figure S2). Therefore, RNAseq data was further explored to assess whether expression of nuclear hormone receptors is significantly altered in ferroptotic MM1 cells. Supplementary Figure S3 demonstrates that most hormone receptors, including estrogen (ESR), glucocorticoid (NR3C1), retinoid X (RXR), and peroxisome proliferator-activated receptors (PPAR) remain largely unaltered upon RSL3 treatment. Similarly, we could not detect significant differences in ferroptosis sensitivity in glucocorticoid-sensitive MM1S cells, expressing NR3C1, versus glucocorticoid-resistant MM1R cells, lacking functional NR3C1 expression. In contrast, an increase in mRNA expression of lipid and oxidative metabolism sensing orphan nuclear receptors NR4A1, NR4A2, and NR4A3 could be observed in RSL3-treated cells compared to untreated controls, and was partly validated by Western blot (Supplementary Figure S4). The interplay between FOXA1 and orphan nuclear receptors has only poorly been characterized, mostly in context of dopaminergic neurons [45, 46]. Interestingly, important tumor suppressor roles for NR4A TFs have recently been described (reviewed in [47]), and NR4A defects are reported to promote formation of blood-tumors (e.g. leukemia, lymphoma) and T-cell immunity dysfunctions [48-51]. As such, possible anti-tumor functions of NR4A TFs in ferroptotic cells deserves further investigation.

We next explored whether FOXA1 might play a role in epithelial-to-mesenchymal transition (EMT) as reported in previous studies [40, 41, 52]. Correlation analysis of the RNAseq data indeed demonstrated that genes highly correlated with FOXA1 expression were enriched in cytoskeleton organization, epithelial cell differentiation, and regulation of EMT (Figure 3a-c). Interestingly, the EMT status is known to directly affect ferroptosis sensitivity, with mesenchymal cells being more susceptible to ferroptotic cell death compared to epithelial cells [43]. Ferroptosis-mediated upregulation of FOXA1 might subsequently drive MM1 cells towards a mesenchymal profile and promote cell death by RSL3. To this end, qPCR analysis of four key EMT markers was performed on MM1 cells treated with RSL3 for increasing timepoints (Supplementary Figure S5). Overall, no significant expression differences of epithelial marker E-cadherin (E-CAD) or mesenchymal markers N-cadherin (N-CAD), Twist-related protein 1 (TWIST1) or Snail Family transcriptional repressor 2 (SLUG) could be observed in ferroptotic cells. These preliminary results indicate that FOXA1 does not orchestrate trans-differentiation of MM1 cells into a mesenchymal phenotype.

**Figure 3:**
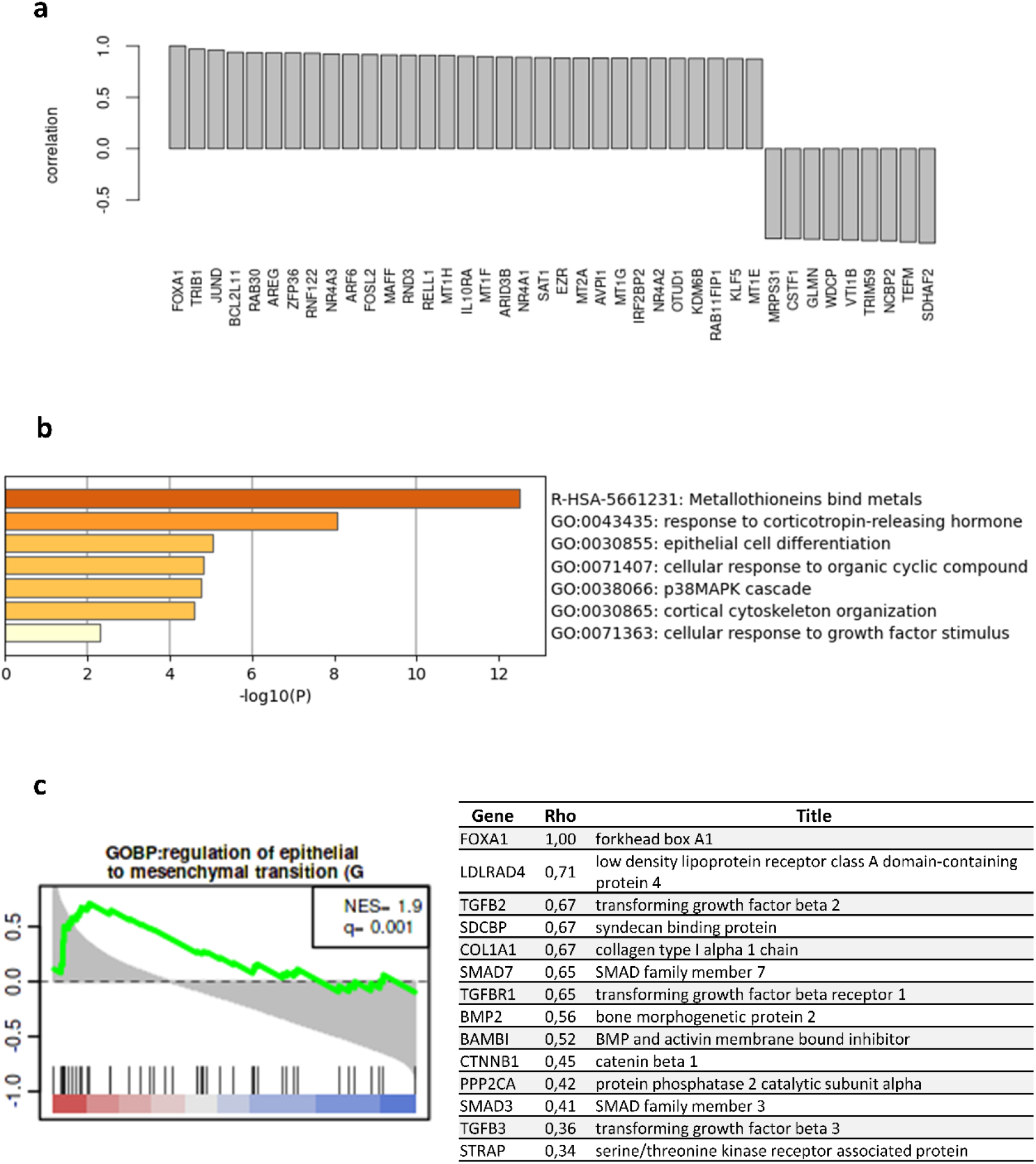
(**a**) Histogram plot displaying the top correlated genes in respect to FOXA1. The height of the bars corresponds to the Pearson correlation value. Figure was generated using the Omics Playground tool (v2.7.18). (**b**) Metascape pathway analysis [57] of RNAseq data displaying the top 20 significantly enriched pathways of RSL3-treated MM1 cells compared to untreated controls. (**c**) Functional GSEA enrichment of genes correlated with FOXA1 expression (left). The green curve corresponds to the normalized enrichment score (NES). Black vertical bars indicate the rank of genes in the gene set in the sorted correlation metric. FDR is represented by the q-value in the figure. Figure was generated using the Omics Playground tool (v2.7.18). The leading-edge table (right) reports the leading edge genes as reported by GSEA corresponding to the selected geneset. The ‘Rho’ columns report the correlation with respect to FOXA1.

Since our targeted approaches did not further elucidate the role of FOXA1 in ferroptosis signaling, we aimed to characterize the downstream effects of FOXA1 by performing CUT&RUN sequencing. This technique allows for genome-wide profiling of chromatin binding sites of transcription factors, similar to ChIP-Seq [53]. In short, MM1R cells were treated for 3 hours with 5 µM RSL3, after which FOXA1-bound DNA fragments were collected and purified for downstream analysis. After completing library preparation and DNA sequencing, enriched regions were called using the sparse enrichment analysis for CUT&RUN (SEARC) [54]. Only a limited number of genomic regions (n = 43) were identified to be differentially altered in FOXA1 binding after RSL3 treatment (Supplementary Table S2). Although we previously measured higher FOXA1 expression in ferroptotic cells, untreated controls displayed higher FOXA1 binding compared to RSL3-treated cells (Supplementary Table S2). Remarkably, all identified regions were located in pericentromeric DNA (Figure 4a-b), suggesting that a ferroptosis-mediated loss of FOXA1 binding to pericentromeric hetero-chromatin takes place upon RSL3 induction. Given that FOXA1 is a pioneer TF that is able to bind compact DNA, these observations could indicate that DNA decondensation (of pericentromeric regions) triggers genome-wide loss of FOXA1 binding. In line with these results, we previously found (unpublished data) that ferroptosis might induce early cellular senescence in MM1 cells, a process which has been associated with defective pericentric silencing and decondensation [55, 56]. Preliminary Western blot analysis of RSL3-treated MM1R cells confirmed the observed loss of FOXA1 expression in chromatin-bound cellular protein fractions (Figure 5). In parallel, cytoplasmic protein expression of FOXA1 was slightly increased upon RSL3 exposure, indicating that chromatin-free FOXA1 is transported toward the cytoplasmic compartment (Figure 5).

**Figure 4:**
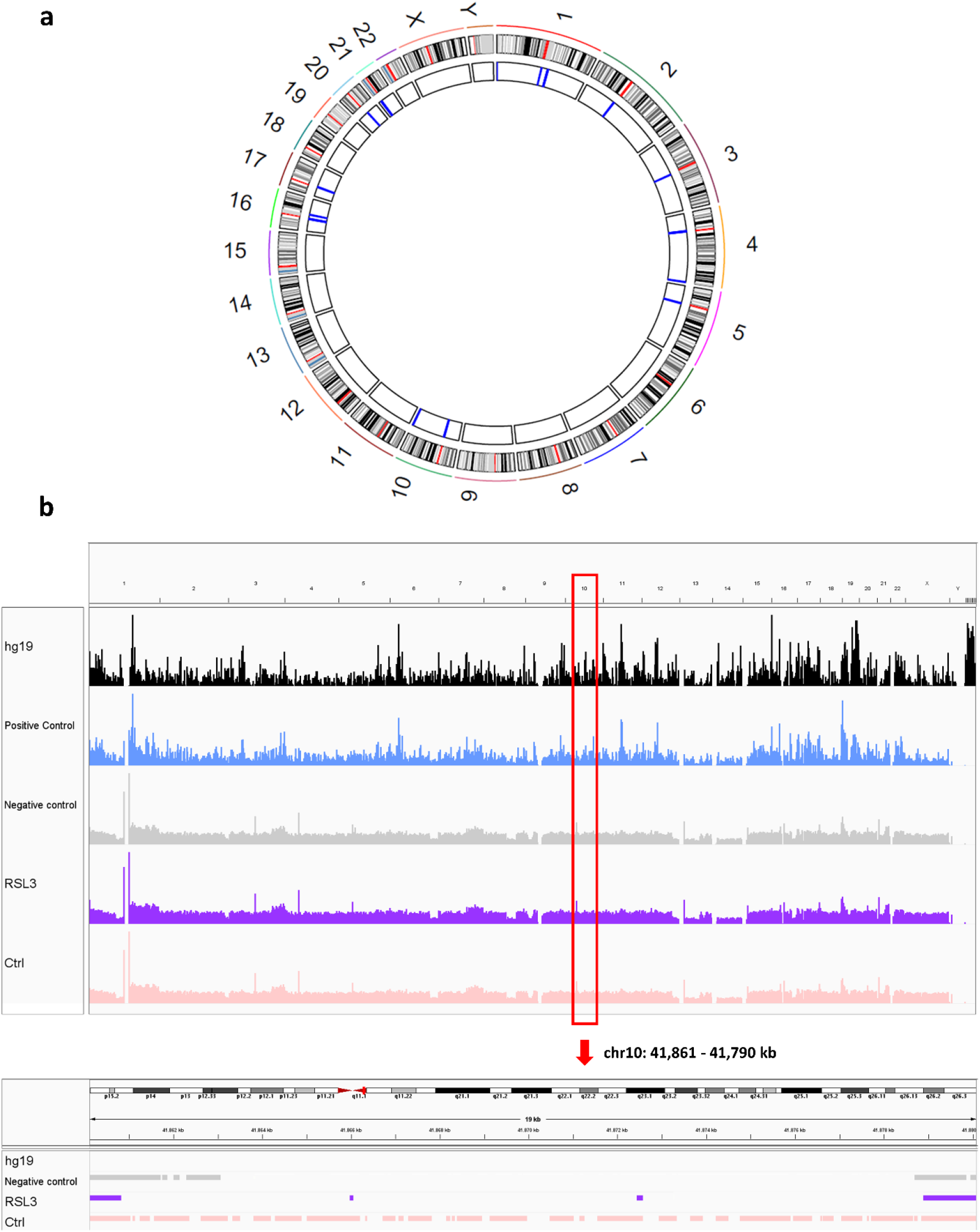
(a) Circos plot displaying chromosome ideograms (outer ring) with centromeric regions marked in red. Blue markings on the inner ring show the differentially enriched FOXA1-bound DNA regions in untreated control cells compared to RSL3-treated cells. (b) Overview of genomic regions sequenced in each CUT&RUN treatment condition (upper panel). The hg19 panel represents the reference genome and displays known mapped genes. The lower panel represents a close-up visualization of chr10: 41,861 – 41,790 kb highlighting the loss of FOXA1 binding to perichromatin in RSL3-treated cells compared to controls. Figures were generated with Integrative Genomics Viewer (v2.9.4).

**Figure 5:**
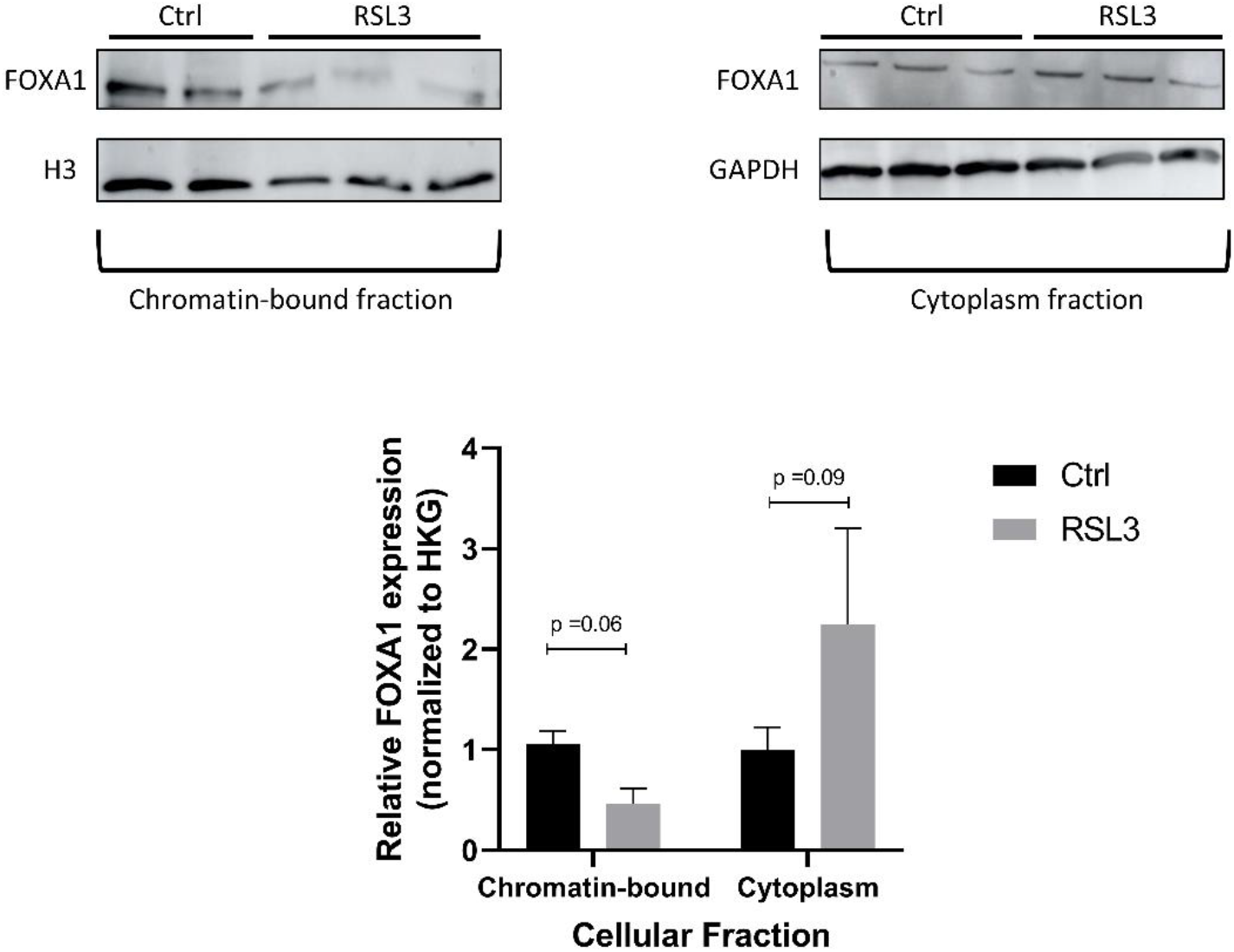
Relative FOXA1 protein expression of different cellular protein fractions in MM1R cells treated with 5 µM RSL3 compared to untreated controls. Data are plotted as the mean ± s.d., *n*= 3 independent samples per treatment (indicated p-values are outcomes of unpaired, two-tailed t-test).

### Sp1 is a Possible Driver of FOXA1 Expression

Taking into account that FOXA1 is upregulated under different ferroptotic conditions in different cell lines, we investigated whether a common transcription factor drives expression of FOXA1. To this end, we generated a list of potential FOXA1 driver genes by identifying transcription factor binding sites located in the FOXA1 promotor region obtained from the SwissRegulon database [58]. Next, a target list of each of the candidate drivers was constructed using three different databases, namely IFTP, TRRUSR, and Marbach2016, employed in the tftarget R package [59]. Finally, an overlap between candidate driver target genes and significant DEGs identified in the two RNAseq studies was performed. Our analysis showed that transcription factor Sp1 is the most probable driver in FOXA1 expression, both in MM1R cells and HEPG2 cells (Table 2).

**Table 2:** Top 5 candidate drivers of FOXA1 expression in ferroptotic cells.

| Candidate driver | # DEGs in RNAseq data regulated by candidate driver | % DEGs in RNAseq regulated by candidate driver |
| --- | --- | --- |
| Sp1 | 46 | 82.14 |
| SPI1 | 36 | 64.29 |
| TFAP2A | 33 | 58.93 |
| TFAP2C | 31 | 55.36 |
| RREB1 | 30 | 55.57 |

Abbreviations: Sp1, Sp1 transcription factor; SPI1, Spi-1 proto-oncogene; TFAP2A, Transcription factor AP-2 alpha; TFAP2C, Transcription factor AP-2 gamma; RREB1, Ras responsive element binding protein 1

Although other studies have reported a ferroptosis-dependent increase in Sp1 expression [7, 60, 61], we did not find significant alterations in Sp1 mRNA levels in RSL3-treated MM1 cells compared to untreated controls (Figure 6a). Nonetheless, preliminary experiments demonstrate that siRNA silencing of Sp1 abolishes ferroptosis-driven FOXA1 upregulation (Figure 2a, Figure 6b), suggesting that Sp1 (partly) drives FOXA1 upregulation in MM1 cells.

**Figure 6:**
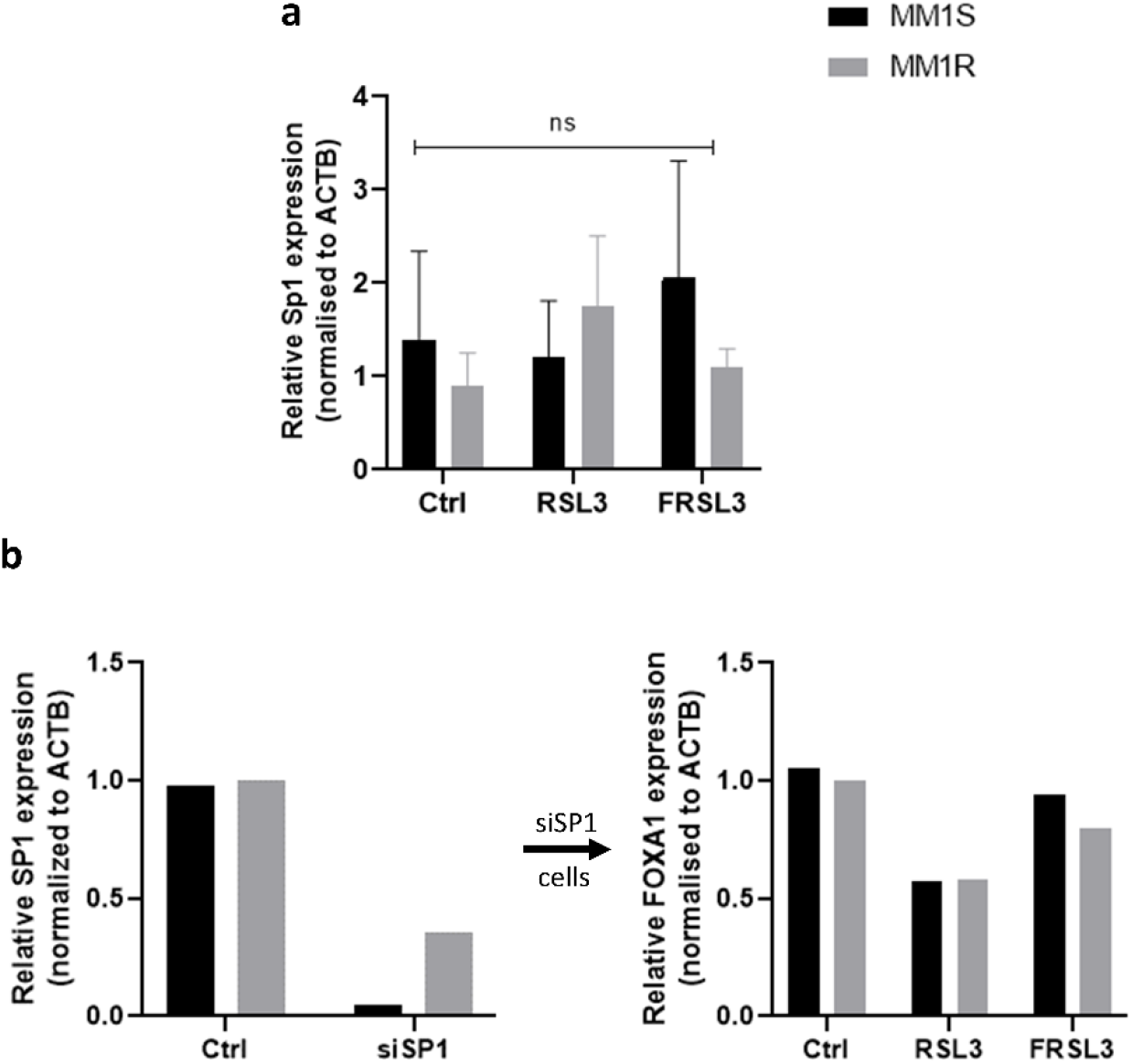
(**a**) Relative mRNA Sp1 expression in MM1 cells after 3 hr treatment with 5 µM RSL3, with (FRSL3) or without pre-treatment with ferrostatin. Data are plotted as the mean ± s.d., *n*=3 biologically independent samples per cell line (ns = p > 0.05, ANOVA). (**b**) Relative mRNA SP1 (left) and FOXA1 (right) expression in MM1 cells transfected with siRNA targeting SP1. Data are plotted as the mean ± s.d., *n=*1 biological independent sample per cell line

Possibly, ferroptotic triggers regulate Sp1 transcriptional activity through alternative mechanisms and subsequently promote downstream FOXA1 expression. In agreement with this hypothesis, a recent kinome screen has revealed that Sp-1 upstream ATR damage response serine/threonine kinase directly impacts ferroptosis sensitivity [62]. Sp1 phosphorylation is known to directly impact Sp-1 dependent transcription [63] and might be increased in ferroptotic cells. Alternatively, ferroptosis signaling pathways could increase expression of Sp1 cofactors and promote Sp1 target site binding. NR4A1 is a cofactor of Sp1 that has recently been described in ferroptotic cell death [64] and was also found to be upregulated in this study. Through its interaction with Sp1, NR4A1 might recruit Sp1 more efficiently to its GC-ich gene targets and promote transcription [65]. Further research exploring the Sp1-FOXA1 signaling axis during ferroptosis are needed to fully confirm the role of Sp1 in RSL3-dependent FOXA1 expression.

## Discussion

In the past decade, RCD and ferroptosis research has grown rapidly, especially in the field of neurological diseases and oncology [66]. Several morphological, biochemical, genetic, and protein hallmarks of ferroptotic cell death have been identified over the last years, but the exact executioner signals of ferroptosis remain largely unknown (reviewed in [29]). Identification of specific ferroptosis contributors may therefore provide novel opportunities for creating anti-cancer therapies. Consequently, we compared RNAseq data of ferroptotic MM and HEPG2 cells to explore whether common expression signatures could be found in these different experimental setups. Our analysis revealed that genes involved in metal binding, DNA binding, protein ubiquitination, and protein phosphorylation were shared in both ferroptosis models. Of particular interest, we identified ATP-independent chromatin remodeler FOXA1 to be specifically upregulated upon RSL3 and erastin treatment. FOXA1 levels were also found to be upregulated in liver tissue obtained from liver-specific GPX4 inducible knock-out mice, suggesting that increased FOXA1 mRNA and protein expression might be a universal trigger in various ferroptosis disease models. Further *in silico* motif and TF enrichment analysis predicted that Sp1 is the most likely driver of ferroptosis-driven FOXA1 expression. Sp1 has previously been described in context of lipid peroxidation and ferroptotic cell death, and is hypothesized to play a dual role in the regulation of tissue injury [7, 60, 61]. However, our qPCR data demonstrate that Sp1 expression remains unaltered in RSL3-treated MM1 cells, implying that transcriptional activity of Sp1 is orchestrated through other upstream mechanisms. Post-translational modifications (PTMs), such as protein phosphorylation, are reported to directly influence Sp1 activity and might be altered in ferroptotic conditions [67]. Indeed, several upstream kinases responsible for Sp1 phosphorylation, including p38 and ATM/ATR are known to be involved in ferroptosis signaling as well [62, 68-70]. Alternatively, transcription activity of Sp1 may be stimulated through improved recruitment to its DNA target binding sites by cofactor proteins that are differentially expressed in the presence of ferroptotic stimuli. NR4A1, for example, has recently been identified as a modulator of ferroptotic cell death and is also reported to act as a cofactor of Sp1 [64, 65]. Follow-up proteomics, Western blot, and immunoprecipitation experiments will undoubtedly reveal to which extend PTMs and cofactor-recruitment of Sp1 are crucial for FOXA1 expression.

Two independent studies have recently reported that nuclear hormone receptor activity is highly correlated with ferroptosis sensitivity [71, 72]. Presumably, cells with higher endocrine activity are subjected to hormone-dependent ROS production, which promotes lipid peroxidation through activation of Fenton reactions [73]. Because FOXA1 is a critical interacting partner of several nuclear receptors [74], we wondered whether ferroptosis induction in MM1 cells is associated with an increase in hormone receptor activity. A preliminary screening of hormone receptor expression mRNA changes in RSL3-treated MM1 cells showed that the expression of the majority of nuclear receptors remain unchanged. Only a subset of orphan nuclear receptors, NR4A1-3, are specifically upregulated upon RSL3 induction. Although FOXA1 has been reported to regulate NR4A2 expression in immature midbrain dopaminergic neurons [45], the interplay between both proteins needs to be explored further. Possibly, NR4A receptors mediate ferroptotic cell death by influencing the cellular energy and lipid metabolism [64, 75]. On the other hand, these orphan receptors might orchestrate ferroptosis signaling pathways by recruiting other ferroptosis-dependent proteins to their target site, as previously explained. NR4A orphan receptors have also been associated with tumor suppressor functions, and mutations in NR4A1-3 have been linked with the formation of blood cancers, including leukemia and lymphoma [48, 49]. RSL3-driven upregulation of NR4A TFs might therefore drive elimination of MM cancer cells by regulating key cancer pathways (reviewed in [47]. Intriguingly, both ferroptosis and NR4A proteins are known to be regulated by the p53 tumor suppressor [76, 77] and indicates that an GPX4-NR4A-p53 signaling network may drive MM cell death. Further research about the anti-cancer effects of NR4A TFs in ferroptotic cells could potentially offer new therapeutical insights for MM and other hematological malignancies. Regardless, based on assessing expression changes, FOXA1 does not seem to primarily target (steroid) hormone receptors during ferroptosis. A direct measurement of hormone receptor activity, by evaluating nuclear translocation or by performing ChIP for example, might aid in fully characterizing the effects of FOXA1 in hormone signaling. Furthermore, evaluating FOXA1-dependent changes on nuclear receptors in more endocrine active cell systems, such as breast or pancreas cancer cell lines, might reveal cell type-dependent effects of FOXA1.

Because both FOXA1 and ferroptosis have been associated with EMT [40, 43, 52], we also investigated whether RSL3 treatment triggers significant changes in EMT markers. Generally, (tumor) cells harboring a more mesenchymal profile are considered to portray an increased ferroptosis sensitivity because they heavily rely on GPX4 activity compared to their epithelial counterparts [78]. Mesenchymal-state cells also exhibit more dysregulated antioxidant programs, explaining why ferroptotic compounds are more potent in these cells [78, 79]. In this regard, ferroptosis-dependent reprogramming of the epithelial-mesenchymal state through FOXA1 upregulation, might promote ferroptosis sensitivity. While our RNAseq data revealed a correlation of FOXA1 expression with several other drivers of EMT, including TRIB1, N-CAD, E-CAD, SLUG, and TWIST1 mRNA expression was not significantly altered upon RSL3 incubation. This suggests that MM1 cells do not shift toward a more epithelial – or mesenchymal-like state under ferroptotic conditions.

Given that neither hormone signaling or EMT seem to be direct downstream targets of FOXA1 in ferroptotic MM1 cells, genome-wide transcription site profiling was performed by CUT&RUN. Similar to ChIP-Seq, this technique combines ChIP with parallel DNA sequencing to identify binding sites of DNA-associated proteins, such as FOXA1. Unfortunately, signal-to-noise signals were quite low in our treatment setups, with signal intensities being similar to the negative IgG control. Further optimization of the experimental setup is therefore required before biologically relevant interpretations can be finalized. Increasing the starting amount of MM1 cells or addition of an extra cross-linking step might improve experimental outcome, especially since FOXA1 has been reported to transiently bind to its DNA sites [80]. Taking this into account, we could still identify 43 genome regions wherein FOXA1 binding was significantly altered in RSL3-treated MM1 cells compared to their untreated controls. Remarkably, all these regions were located in pericentromeric chromatin and FOXA1 binding was significantly lower in RSL3 conditions, despite the earlier observed transcriptional and translational FOXA1 upregulation in MM1 cells. Pericentric (satellite) DNA is typically considered to be void of functional genes and transcriptionally silent since they are confined in transcriptionally inert heterochromatin [81]. However, mounting evidence suggests that pericentric transcripts are crucial in maintaining genome stability (reviewed in [82] and [83]). To this end, loss of FOXA1 binding in ferroptotic cells could potentially promote genome instability and DNA double strand breakage. Another possibility is that ferroptotic stress triggers defective pericentric transcription due to dramatic DNA decondensation, as is also observed when cells are exposed to UV (i.e. DNA damage), cadmium toxicity or cellular senescence [56, 84-86]. Given that FOXA1, as a pioneer TF, mainly binds to heterochromatin regions, genome-wide DNA decondensation might promote overall loss of FOXA1 pericentromeric DNA binding and uncontrolled pericentric transcription. Our previous work (unpublished data) has indeed suggested that ferroptosis is associated with an epigenomic stress response linked to oxidative stress and cellular senescence, suggesting that DNA decondensation might occur in ferroptotic cells. Intriguingly, FOXA1 expression has been reported to increase with cellular senescence [87]. Possibly, loss binding to heterochromatin pericentromeric DNA promotes recruitment of FOXA1 to other target sites that trigger cellular senescence [87]. Repeating the CUT&RUN experiments under optimized experimental conditions should help in investigating this hypothesis further. Alternatively, “DNA-free” FOXA1 might localize to the cytoplasm and inhibit nuclear translocation of other TFs to promote cell death [88]. This seems to occur in our experimental setup as well, given that Western blot analysis revealed an RSL3-dependent increase in FOXA1 expression in cytoplasmic protein fractions.

Taken together, our data suggest that ferroptosis triggers a time-dependent upregulation of FOXA1 expression in different experimental models, which could be orchestrated by transcriptional activation of Sp1. The downstream effects of this FOXA1 expression surge in MM1 cells remain somewhat elusive but do not seem to include steroid hormone signaling or EMT. In contrast, preliminary data imply that ferroptotic stress might trigger uncontrolled pericentric transcription and genome instability, due to loss of FOXA1-binding to pericentromeric DNA. Moreover, relocalization of pericentric-free FOXA1 to secondary target sites or the cytoplasm might further promote cellular stress responses, such as cellular senescence or cell death.

## Materials and Methods

### Cell Culture and Cell Viability Assays

Human MM1S cells (CRL-2974) and MM1R cells (CRL-2975) were purchased from ATCC. RPMI-1640 medium, supplemented with 10% FBS (E.U Approved; South American Origin) and 1% Pen-Strep solution (Invitrogen, Carlsbad, CA, USA), was used to sustain the cells. The cells were cultivated at 37°C in 5% CO2 and 95% air atmosphere and 95-98% humidity. To assess cell viability, the colorimetric assay with 3-(4, 5-dimethylthiozol-2-yl)-2, 5-diphenyltetrazolium bromide (MTT) was used (Sigma Aldrich, St. Louis, MO, US) as previously described [89].

### Antibodies and Reagents

RSL3 was purchased from Selleckchem (Houston, USA), dissolved in DMSO and stored as 50 mM stocks at -20°C. siRNA targeting Sp1 (1299001) was purchased from ThermoFisher Scientific (Waltham, MA, USA)). Antibodies FOXA1 (ab23738) and GAPDH (2118S) were obtained from Abcam (Cambridge, UK) and Cell Signaling Technology (Danvers, MA, USA), respectively.

### RNA Extraction and Sequencing

After cell harvest, total RNA from untreated or RSL3-treated (with or without 2 hr pre-treatment with 2 µM Ferrostatin-1) MM1S and MM1R cells was extracted using the RNeasy Mini Kit (Qiagen, Venlo, the Netherlands) according to the manufacturer’s protocol. Once isolated, quantification was performed with the Qubit RNA BR Assay Kit (ThermoFisher, MA, USA) and RNA was stored at -80°C. Extracted RNA was used as input for RNA sequencing as previously described [90]. In brief, RNA was shipped to BGI (BGI Group, Bejing, China) where quality checks were performed using the 2100 Bioanalyzer system (Agilent Technologies, USA) and sequencing took place using the BGISE-500 platform (BGI Group, Beijing, China). Quality control, genome mapping and differential gene expression analysis was performed using the R-packages FastQC (v0.11.5) [91], STAR (v2.7.3a) [92], and DESeq2 (v3.12) [93]. DEGs were considered to be significant when FDR < 0.05 and │log2FC│> 1. Raw gene counts from the GSE104462 dataset [94] were extracted from the Gene Expression Omnibus (GEO) database and used as input for the same RNAseq analysis pipeline as described above.

### cDNA Synthesis and Quantitative Real-time PCR

Extracted RNA from RSL3-treated cells was converted into cDNA using the Go-Script reverse transcription system (Promega, Madison, Wisconsin, USA) according to the manufacturer’s protocol. Subsequently, qPCR analysis was carried out using the GoTag qPCR Master Mix (Promega, Madison, Wisconsin, USA) as explained by manufacturer’s protocol. In short, 1µL cDNA was added to a master mix comprising SYBR green, nuclease-free water, and 0.4 µM forward and reverse primers. The following PCR program was applied on the Rotor-Gene Q qPCR machine (Qiagen, Venlo, the Netherlands): 95°C for 2 min, 40 cycli denaturation (95°C, 15 s) and annealing/extension (60°C, 30 s), and dissociation (60–95°C). Each sample was run in triplicate and the median value was used to determine the ΔΔCt-values using β-actin (BACT) as the normalization gene. Primer sequences are listed in Supplementary Table S1.

### Protein Extraction and Western blot Analysis

Cellular protein extraction occurred by resuspending cell pellets in 0.5 mL RIPA buffer (150 mM NaCl, 0.1% Triton X-100, 1% SDS, 50 mM Tris-HCl pH 8) supplemented with PhosphataseArrest (G-Biosciences, Saint-Louis, MO, USA) and protease inhibitors (Complete Mini®, Roche). After 15 min incubation on ice with regular vortexing, samples were briefly sonicated (1 min, amplitude 30 kHz, pulse 1s) and centrifuged at 13 200 rpm for 20 min at 4°C. Solubilized proteins were transferred to new Eppendorf tubes and stored at -20 °C. To extract proteins from mouse liver tissue, RIPA buffer containing 2% SDS was added to the tissue. Sample homogenisation was performed with the TissueRuptor, followed by 1 hour incubation at 4°C on a rotor. Sonication (5 min, low amplitude 1 kHz and 20 Hz burst rate) was used to shear DNA and debris was removed by centrifugation for 8 min at 13 000 g.

Using standard protocols, all protein samples were separated using Bis-Tris SDS-PAGE with a high-MW MOPS running buffer, and transferred onto nitrocellulose membranes (Hybond C, Amersham) using the Power Blotter System (Thermofisher, MA, USA). Blocking the membranes for 1 hour with blocking buffer (20 mM Tris-HCl, 140 mM NaCl, 5% BSA, pH 7,5) at RT was followed by overnight incubation with the primary antibody at 4°C. Blots were then incubated for 1 hr with the secondary, HRP dye-conjugated antibody (Dako, Glostrup, Denmark) after which chemiluminescent signals were detected with the Amersham Imager 680 (Cytiva, MA, USA) and quantified with the ImageJ software (v1.53j) [95].

### Liver Samples of Cre-lox Liver-Specific GPX4 Knockout Mice

Liver samples from Cre-lox liver-specific inducible GPX4 kockout mice were kindly provided by Ines Goetschalckx and Prof. Dr. Tom Vanden Berghe (Laboratory of Pathophysiology, University of Antwerp). GPX4 knockout mice were generated by crossing homozygous GPX4-floxed mice with heterozygous GPX4 conditional knockout mice (Supplementary Figure S1).

### Nucleofection of MM cells

MM1 cells were transfected using the Nucleofector IIb device (Lonza Amaxa, Switzerland) as described by the manufacturer’s protocol. Briefly, 1 million cells were resuspended in 100 µL supplemented nucleofector solution. Next, 300 nM siSp1 was added to resuspended cells. To assess transfection efficiency, an additional pmaxGFP Vector (Lonza, Bazel, Switserland) was included in each nucleofection reaction (average transfection efficiency = 51.3 ± 2.4 %). Cell suspensions were transferred to a provided cuvettes and nucleofection was performed using the O-020 program. 48 hours after transfection, cells were harvested and used as input for qPCR analysis.

### Motif Analysis and Transcription Factor Enrichment Analysis

To search for a potential driver of FOXA1, a list of candidate drivers was composed using transcription factor binding sites from the SwissRegulon database located at the *FOXA1* promoter [58, 96]. Using that list, a list of targets of these candidate drivers was composed using data from three different databases – IFTP, TRRUST and Marbach2016 – provided via the tftargets package in R [59]. Targets of each candidate driver were matched to DEGs common to both datasets and two metrics for overlap were calculated: the number of overlapping genes and the percentage of overlap with respect to the number of DEGs. As an additional control, the X2Kweb tool was further used to identify putative enriched transcription factors through Transcription factor enrichment analysis (TFEA) [97]. Results from TFEA were compared with results from IFTP, TRRUST and Marbach2016.

### CUTANA CUT&RUN to Identify Chromatin-Associated Proteins

Downstream targets of FOXA1 were identified using the EpiCypher CUTANA ChIC CUT&RUN Kit (23614-1048, EpiCypher, USA) as previously described [53, 98]. In short, 5 × 10^5^ MM1R were plated into 6-well plates and either treated with 5 µM RSL3 (3 hours) or left untreated. For each treatment condition, 4 biological replicates were included. Cells were washed and bound to concanavalin A-coated magnetic beads and permeabilized with wash buffer (20 mM HEPES pH 7.5, 150 mM NaCl, 0,5 mM spermidine and protease inhibitors) supplemented with 0,05% digitonin. After overnight incubation at 4°C with the primary FOXA1 antibody (13-2001, Epicypher), cell-bead slurry was washed twice more after which pA-MNase digestion was activated by placing samples on an ice-cold block and incubated with digitonin wash buffer containing 2 mM CaCl2. Each CUT&RUN experiment also featured a positive (anti-H3K4me3) and negative (anti-Rabbit IgG) antibody control. After 2 hours, the cleavage reaction was stopped with stop buffer (340 mM NaCl, 20 mM EDTA, 4 mM EGTA, 0.05% digitonin, 0.05 mg/mL glycogen, 5 µg/mL RNase A, 2 pg/mL *E. coli* spike-in DNA) and fragments were released by 30-minute incubation at 37°C. Samples were centrifuged and DNA-containing supernatant was collected. DNA extraction was performed with a DNA extraction kit supplied with the CUTANA Kit. Resulting DNA was used as input for library preparation using Kapa HypePrep Kit (7962363001, Roche) and barcoding using xGen UDI-UMI barcodes (10005903, IDT) following manufacturers protocols. Barcoded libraries of ten samples (4 treated, 4 untreated and two controls) were equimolarly pooled and sequenced on MiSeq (Illumina) using MiSeq v3 150 reagent kit (MS-102-3001, Illumina) according to manufacturer’s protocol. The run ended in obtaining 4.16 Gb, 56.66 million reads (Q30 92.09%, 3,86 Gb).

Sequencing data were aligned to the UCSC h38 reference genome using the Burrows-Wheeler Aligner [99] and peaks were called using SEARC [54] after conversion to the bedgraph format using bedtools [100]. Peaks were merged using Granges [101]. For each peak region, the number of mapping reads was counted using chromVAR [102] getCounts. *E*.*coli* spike-in counts were obtained by alignment to the eschColi_K12 *E*.*coli* reference. Differential analysis was performed using the DESeq2 package (v3.12) [93], where counts were normalized to *E*.*coli* spike-in counts. Differentially enriched regions were visualized with the RCircos R package (v1.2.1) [103] and IGV (v2.9.4) (BroadInstitute, Cambridge, MA, USA).

### Subcellular Protein Fractionation

The subcellular protein fractionation kit (# 78840, Thermofisher, MA, USA) was used to fractionate proteins into nuclear and cytoplasmic fractions according to the manufacturer’s instructions. The yield of obtained chromatin-bound nuclear proteins and cytoplasmic proteins was determined by the BCA method. Finally, 20 µg of protein from each cellular fraction was used to perform SDS-PAGE and Western blot analysis, as described above.

### Statistical Analysis

Statistical tests were performed in GraphPad Prism (v7.0) (GraphPad Software, San Diego, CA, USA) unless otherwise stated in the main text. Results were considered to be statistically significant when p-values < 0.05 were obtained.

