## Supplementary Material for "Investigating the Role of Chromatin Remodeler FOXA1 in Ferroptotic Cell Death"

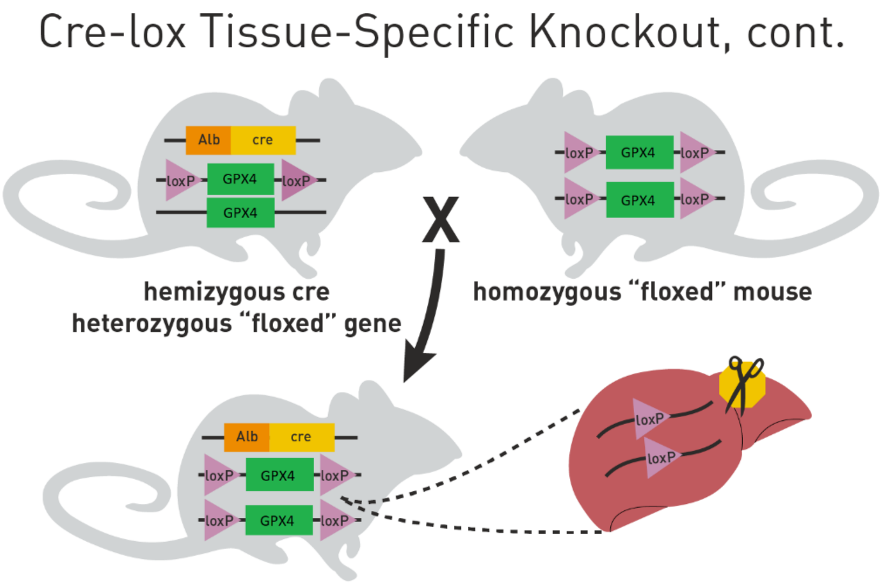


*Figure S1:* Generation of liver-specific, inducible GPX4 knockout mice. Heterozygous GPX4-floxed mice carrying the *cre* transgene are crossed with homozygous GPX4-floxed mice to obtain homozygous loxP-flanked GPX4 mice that are heterozygous for the *cre* transgene. Figure adapted from https://www.jax.org/news-and-insights/jax-blog/2011/september/cre-lox-breeding#.

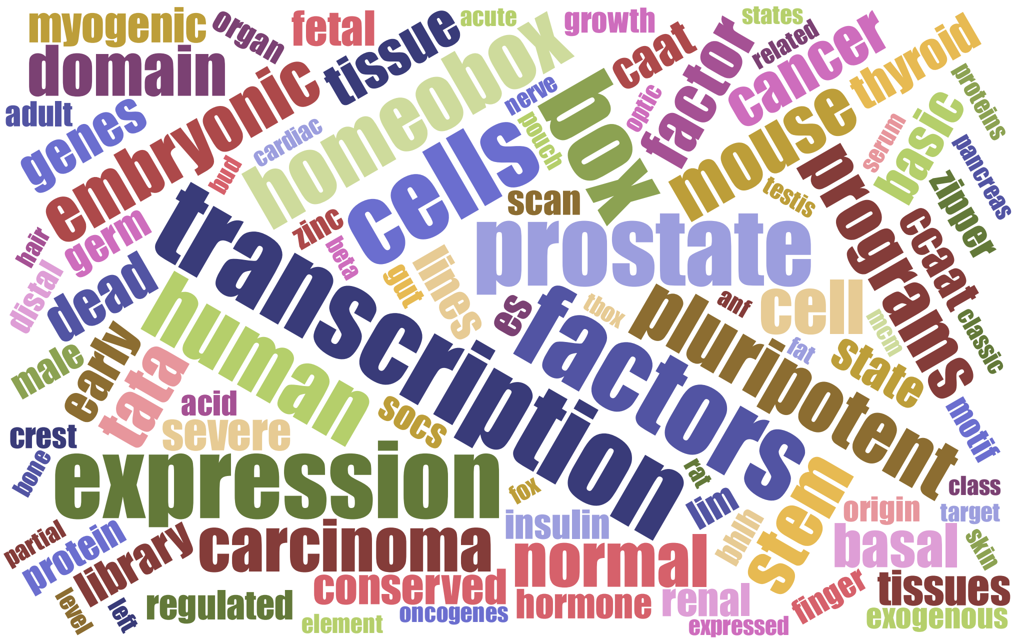


*Figure S2:* Word cloud displaying word associations with FOXA1 obtained from PubMed abstracts and free PubMed full-length articles. The strength of the association is represented by the word size. Word cloud was generated using the online webtools Textrous! and Jason Davies word cloud generator.

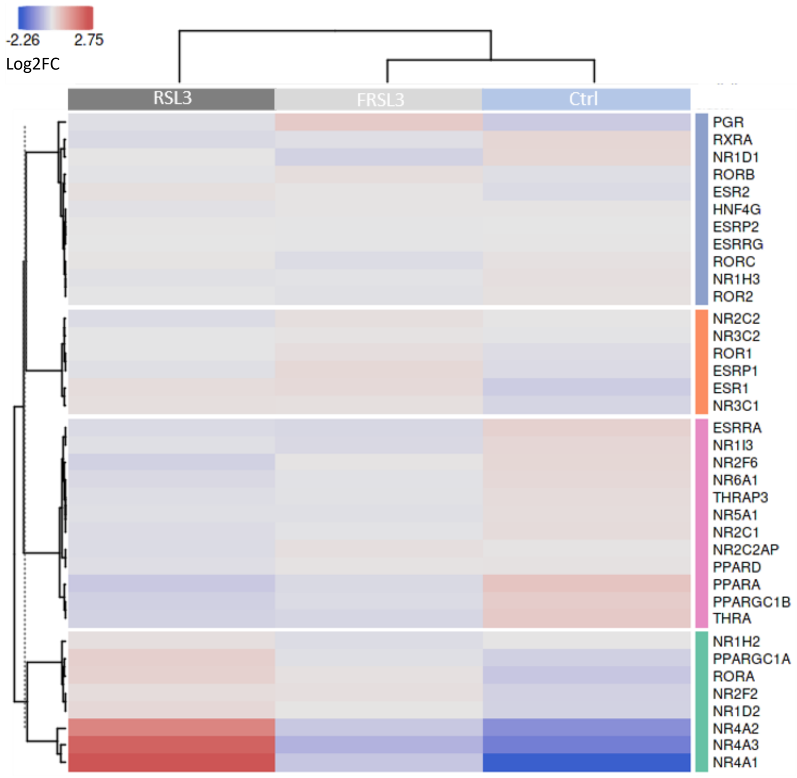


*Figure S3:* Heatmap representation of expression changes (log2FC) of nuclear receptors in RSL3-treated cells (with or without Ferrostatin-1 pre-treatment (FRSL3)) and untreated controls. Heatmap was generated using Omics Playground Tool (v2.7.18).

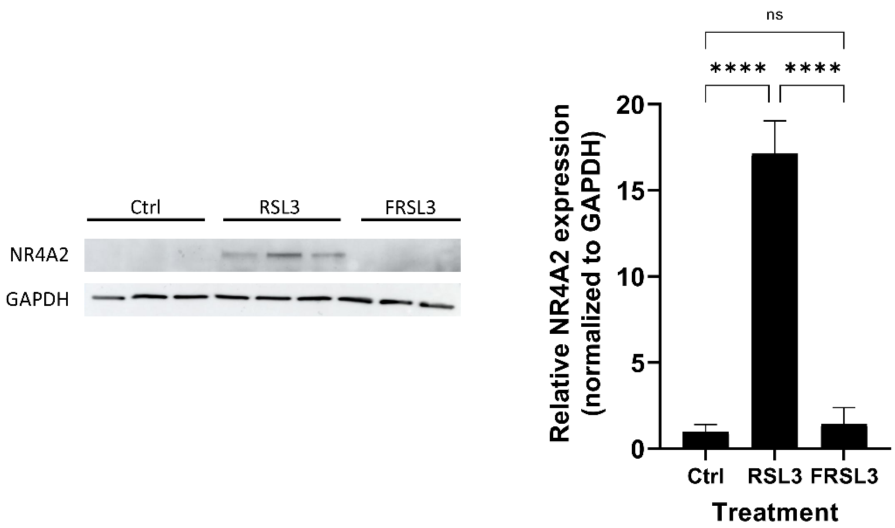


*Figure S4:* Relative NR4A2 protein expression in MM1S cells treated with 5 µM RSL3 (with or without Ferrostatin-1 pre-treatment (FRSL3)) compared to untreated controls. Data are plotted as the mean ± s.d., *n*= 3 independent samples per treatment (ns $p > 0.05$, ****$p < 0.0001$), ANOVA).

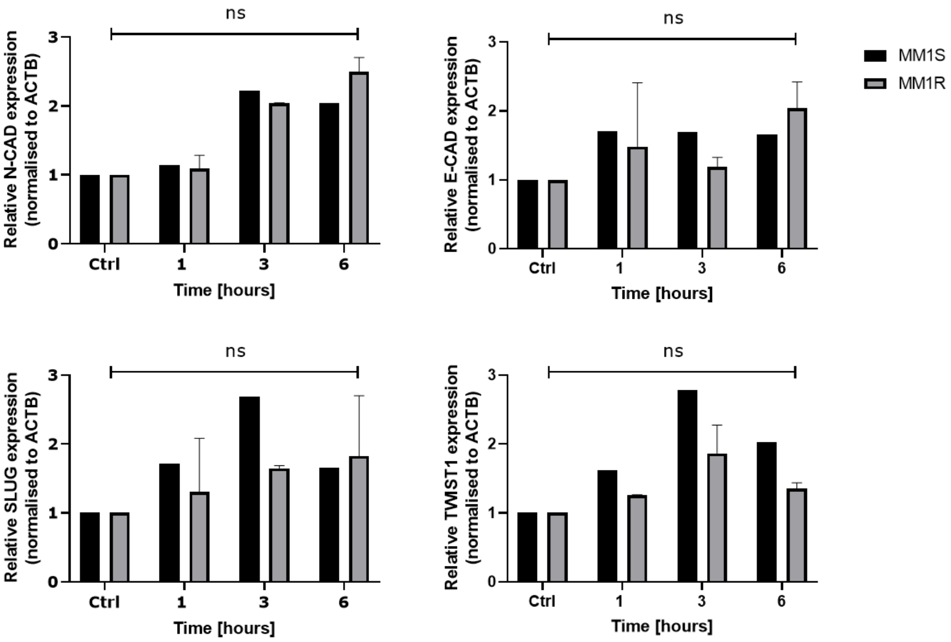


*Figure S5:* Relative mRNA expression of different EMT markers in MM1R and MM1S cells treated with 5 µM RSL3 compared to untreated controls. Data are plotted as the mean ± s.d., *n*=1 or 2 (indicated by presence of error bar) biologically independent samples per cell line (ns $p > 0.05$), ANOVA). Abbreviations: N-CAD = N-cadherin, E-CAD = E-cadherin, SLUG = snail family transcriptional repressor 2, TWIST1 = twist family BHLH transcription factor 1.

*Table S1:* Overview of primers used in this study

| Target | | Primer Sequence | Gene accession number |
|---|---|---|---|
| *qPCR primers* | | | |
| FOXA1 | Forward<br>Reverse | AGG TGT GTA TTC CAG ACC CG<br>TTG ACG GTT TGG TTT GTG TG | ENSG00000129514 |
| Sp1 | Forward<br>Reverse | TTG AAA AAG GAG TTG GTG GC<br>TGC TGG TTC TGT AAG TTG GG | ENSG00000185591 |
| ACTB | Forward<br>Reverse | CTG GAA CGG TGA AGG TGA CA<br>AAG GGA CTTC CTG TAA CAA TGC A | ENSG00000075624 |
| E-CAD | Forward<br>Reverse | TGC CCA GAA AAT GAA AAA GG<br>GTG TAT GTG GCA ATG CGT TC | ENSG00000039068 |
| N-CAD | Forward<br>Reverse | GAC AAT GCC CCT CAA GTG TT<br>TCA CAC GCA GGA TGG AAA TA | ENSG00000170558 |
| SLUG | Forward<br>Reverse | CTT TTT CTT GCC CTC ACT GC<br>GCT TCG GAG TGA AGA AAT GC | ENSG00000019549 |
| TWIST1 | Forward<br>Reverse | AGT CCG CAG TCT TAC GAG GA<br>CAT CTT GGA GTC CAG CTC GT | ENSG00000122691 |

*Table S2:* Overview of significantly enriched genome regions bound to FOXA1 in ferroptotic cells compared to untreated controls

| Chromosome | Start | End | Width | Log2FoldChange | pvalue | padj |
|---|---|---|---|---|---|---|
| chr1 | 125177010 | 125184585 | 7576 | -1,194 | 0,002 | 0,006 |
| chr1 | 143189543 | 143196164 | 6622 | -1,163 | 0,003 | 0,006 |
| chr1 | 143211365 | 143224006 | 12642 | -1,203 | 0,002 | 0,005 |
| chr1 | 143227041 | 143228185 | 1145 | -1,166 | 0,003 | 0,006 |
| chr1 | 143249564 | 143250811 | 1248 | -1,335 | 0,004 | 0,006 |
| chr1 | 143261455 | 143267803 | 6349 | -1,206 | 0,002 | 0,005 |
| chr1_KI270709v1_random | 3587 | 8994 | 5408 | -1,167 | 0,002 | 0,005 |
| chr2 | 89830889 | 89834065 | 3177 | -1,050 | 0,003 | 0,006 |
| chr2 | 89835550 | 89837865 | 2316 | -1,337 | 0,009 | 0,012 |
| chr3 | 93470359 | 93470801 | 443 | -1,328 | 0,001 | 0,005 |
| chr4 | 49091387 | 49093922 | 2536 | -1,070 | 0,008 | 0,010 |
| chr4 | 49094532 | 49100149 | 5618 | -1,232 | 0,002 | 0,005 |
| chr4 | 49143075 | 49146670 | 3596 | -1,147 | 0,004 | 0,006 |
| chr4 | 49153138 | 49155368 | 2231 | -1,050 | 0,003 | 0,006 |
| chr4 | 49632314 | 49635881 | 3568 | -1,118 | 0,003 | 0,006 |
| chr4 | 49636565 | 49641731 | 5167 | -1,097 | 0,012 | 0,015 |
| chr4 | 190177041 | 190180141 | 3101 | -1,058 | 0,004 | 0,006 |
| chr5 | 49656292 | 49661874 | 5583 | -1,208 | 0,002 | 0,005 |
| chr5 | 49666169 | 49667431 | 1263 | -1,330 | 0,000 | 0,005 |
| chr10 | 41859110 | 41860822 | 1713 | -1,227 | 0,003 | 0,006 |
| chr10 | 41879171 | 41879681 | 511 | -1,230 | 0,001 | 0,005 |
| chr10 | 41881948 | 41884082 | 2135 | -1,142 | 0,020 | 0,022 |
| chr10 | 133687126 | 133690435 | 3310 | -1,141 | 0,004 | 0,006 |
| chr16 | 34571516 | 34576751 | 5236 | -1,260 | 0,001 | 0,005 |
| chr16 | 34584655 | 34587633 | 2979 | -1,291 | 0,002 | 0,005 |
| chr16 | 34587798 | 34589011 | 1214 | -1,175 | 0,003 | 0,006 |
| chr16 | 34589077 | 34596197 | 7121 | -1,233 | 0,002 | 0,005 |
| chr16 | 46383894 | 46396853 | 12960 | -1,226 | 0,002 | 0,005 |
| chr17 | 26603527 | 26604747 | 1221 | -1,108 | 0,007 | 0,009 |

| | | | | | | |
|---|---|---|---|---|---|---|
| chr17 | 26619202 | 26620327 | 1126 | -1,278 | 0,005 | 0,006 |
| chr17_KI270729v1_random | 2108 | 6136 | 4029 | -1,141 | 0,004 | 0,006 |
| chr17_KI270729v1_random | 20501 | 23628 | 3128 | -1,133 | 0,004 | 0,006 |
| chr20 | 31056797 | 31070093 | 13297 | -1,162 | 0,002 | 0,005 |
| chr21 | 8217544 | 8220689 | 3146 | -0,890 | 0,047 | 0,050 |
| chr21 | 8224769 | 8232425 | 7657 | -1,168 | 0,002 | 0,005 |
| chr21 | 8239285 | 8246018 | 6734 | -1,199 | 0,002 | 0,005 |
| chr21 | 8450219 | 8456017 | 5799 | -1,135 | 0,004 | 0,006 |
| chr21 | 8460222 | 8464400 | 4179 | -1,001 | 0,017 | 0,020 |
| chr22_KI270733v1_random | 133817 | 136973 | 3157 | -0,827 | 0,026 | 0,028 |
| chrUn_GL000216v2 | 5261 | 7418 | 2158 | -1,362 | 0,000 | 0,000 |
| chrUn_KI270333v1 | 93 | 2683 | 2591 | -0,965 | 0,020 | 0,022 |
| chrUn_KI270337v1 | 0 | 1020 | 1021 | -1,049 | 0,013 | 0,016 |
